# Musical experience partially counteracts temporal speech processing deficits in mild cognitive impairment

**DOI:** 10.1101/2021.04.21.440718

**Authors:** Caitlin N. Price, Gavin M. Bidelman

## Abstract

Mild cognitive impairment (MCI) commonly impacts older adults resulting in more rapid cognitive and behavioral declines than typical aging. Individuals with MCI can exhibit impaired receptive speech abilities that may reflect neurophysiological changes in auditory-sensory processing prior to usual cognitive deficits. Benefits from current interventions targeting communication difficulties in MCI are limited. Yet, neuroplasticity associated with musical experience has been implicated in improving neural representations of speech and offsetting age-related declines in perception. Here, we asked whether these experience-dependent effects of musicianship might extend to aberrant aging and offer some degree of cognitive protection against MCI. During a vowel categorization task, we recorded single-channel EEGs in older adults with putative MCI to evaluate speech encoding across subcortical and cortical levels of the auditory system. Critically, listeners varied in their duration of formal musical training experience (0-21 years). Older musicians exhibited sharpened temporal precision in auditory cortical responses suggesting musical experience produces more efficient processing of acoustic features by offsetting age-related neural delays. Additionally, we found robustness of brainstem responses predicted severity of cognitive decline suggesting early speech representations are sensitive to pre-clinical stages of cognitive impairment. Our preliminary results extend prior studies by demonstrating positive benefits of musical experience in older adults with emergent cognitive impairments.

## INTRODUCTION

Cognitive aging results in both structural and functional changes throughout the central auditory pathway^1–3^ as well as declines in associated auditory processing skills necessary for effective speech comprehension^4,5^. Older adults demonstrate less precise and altered neural encoding of speech which has been attributed to slowed neural conduction times, poor neural synchrony, and reduced inhibitory processes^6–8^. Senescent neural changes parallel perceptual difficulties commonly reported by aging adults in clinical settings. Older adults often exhibit deficits in temporal processing which is vital for accurate speech perception particularly in challenging listening conditions such as noisy or reverberant environments^5,9^.

Aging effects are further exacerbated in those with mild cognitive impairment (MCI)^10^, a form of atypical cognitive aging causing memory lapses in short-term recall and/or impaired critical thinking and decision-making skills^11^. Because individuals with MCI are also at higher risk of progressive cognitive deficits with potential conversion from MCI to Alzheimer’s disease or other forms of dementia^11^, early identification and intervention is vital. While most current diagnostic and intervention techniques focus on aspects of cognitive function, recent studies reveal MCI alters early sensory processing by compounding the auditory perceptual deficits due to typical aging^12–14^. Thus, in addition to the more commonly noted cognitive issues, there is emerging evidence that MCI is associated with early declines in *auditory-sensory* processing including complex expressive and receptive language processing^15^ and even how accurately the brain encodes speech itself^12,16^. In this vein, neuroimaging techniques and complex speech perception tasks may aid in the early detection of neurocognitive changes associated with MCI.

Current MCI treatments do little to slow disease progression and primarily address associated symptoms or focus on improving quality of life^11,17^. Interventions targeting communication difficulties in this population provide limited benefit for improving language or cognitive skills^15^. However, studies show musical experience may counteract age-related declines by robustly enhancing the neural encoding of speech at both subcortical and cortical levels of the central auditory system^18–20^. Individuals with musical experience demonstrate stronger cognitive abilities (i.e., working memory and attentional control^21,22^), higher fidelity neural encoding of acoustic features^23,24^, and improved neural timing precision^18,25^, all of which contribute to enhanced speech perception abilities. Nevertheless, while musicianship might help counteract normal age-related declines in speech processing^19,26^, it remains unclear whether such experience-dependent advantages transfer to populations with suspected cognitive impairments. A critical demonstration of music’s benefits on the aging brain is to confirm whether similar neural enhancements occur in at risk populations or those with borderline neurocognitive performance (e.g., MCI). This would provide novel evidence that music engagement might bolster communication skills at early stages and decelerate the progression of cognitive aging.

To this end, the current study aimed to investigate the impact of musical experience on speech encoding in individuals with putative MCI. We measured brainstem frequency-following responses (FFRs) and cortical event-related potential (ERPs) in older adults with varying degrees of musical experience who were identified as being at risk for MCI based on cutoff-level performance on neurocognitive assessments. Our findings reveal that abnormally large brainstem FFRs predict severity of cognitive impairment and may serve as an objective marker for MCI severity. We further show musical experience mitigates some age-related delays in neural speech encoding in this at-risk population, paving the way for future translational music interventions in older populations.

## RESULTS

We evaluated a sample of seventeen older adults (*M±SD*, 71.1 ± 8.0 yrs; range: 52-86 yrs) who demonstrated borderline to slight degrees of cognitive impairment as assessed by the Montreal Cognitive Assessment (MoCA). We separated participants into groups based on their extent of musical experience. Individuals with no prior musical training were included in the nonmusician group (NM; *n =* 8) while those with *any* prior musical training were included in the musician group (M; *n =* 9). Participants performed a categorical perception task in which they were presented with tokens from a synthetic five-step vowel continuum spanning from /u/ to /a/ (hereafter “vw 1-5”). Using a forced choice task, participants identified each vowel token as /u/ or /a/ while we recorded subcortical FFRs and cortical ERPs simultaneously using a single-channel montage.

### Cognitive function

Figure 1 depicts the level of cognitive function for each group as measured by the MoCA. The groups did not differ in MoCA after controlling for age, hearing sensitivity, and education (*F*_1,11_ = 1.24, *p* = 0.29; Fig. 1a). Yet, cognitive function declined with increasing age in both groups (*F*_1,11_ = 8.80, *p =* 0.01, 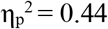; Fig. 1b). Though Ms appeared to have a slower rate of age-related cognitive decline, the group x age interaction did not reach significance (*F*_1,11_ = 1.95, *p =* 0.19).

**Figure 1.**
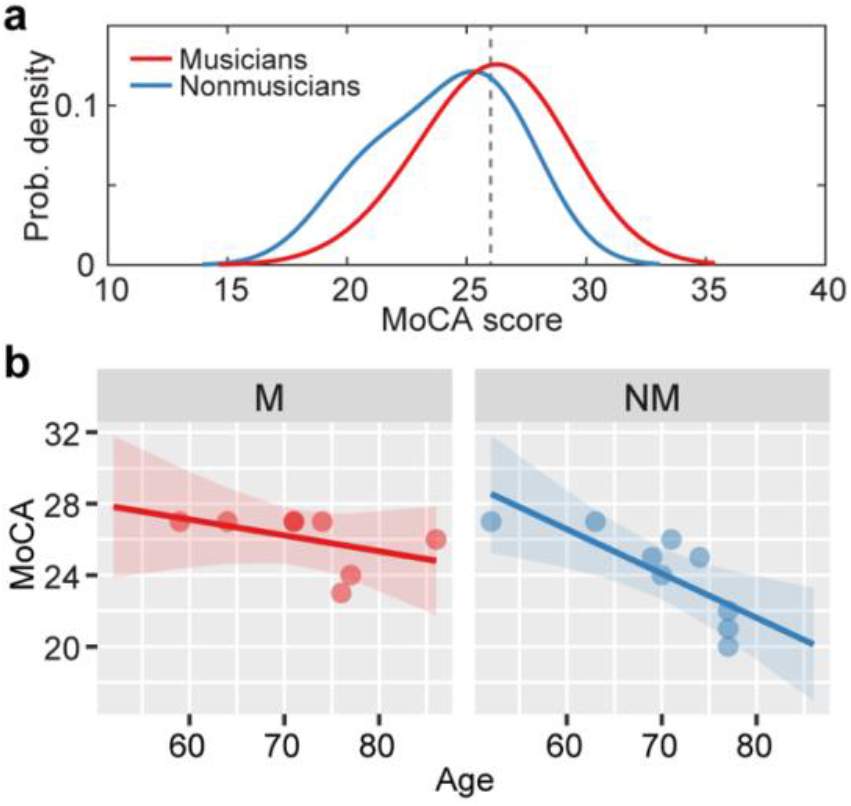
Age-related declines in cognitive function in older musicians and nonmusicians. (a) Distribution of MOCA scores per group. Dotted line, cutoff score for normal cognitive functioning^27^. Older musicians had marginally higher cognitive function than nonmuscians. (b) Cognitive function declines with age for both groups but appears less precipitous in musicians. shading = 95% CI

### Behavioral speech categorization

Ms and NMs demonstrated similar vowel categorization across the continuum (identification function slope: *t*_15_ = −0.90, *p =* 0.38; Fig. 2a). The stair-stepped shape of the curves demonstrate that tokens were perceived categorically as /a/ or /u/, respectively. Likewise, similar patterns in reaction times (RTs) were observed between the groups (*F*_1,60_ = 2.03, *p =* 0.16; Fig. 2b). However, decision speeds varied across vowels (*F*_4,60_ = 15.41, *p <* 0.0001, 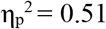). As expected, both groups categorized the prototypical endpoint tokens (i.e., vw 1, 2, 4, 5) more quickly than the ambiguous midpoint token (i.e., vw 3) (M: *t*_23.5_ = −8.97, *p <* 0.001; NM: *t*_23.5_ = −11.32, *p <* 0.001; Fig. 2b), indicating the hallmark slowing of decision speed for category-ambiguous speech sounds^28^.

**Figure 2.**
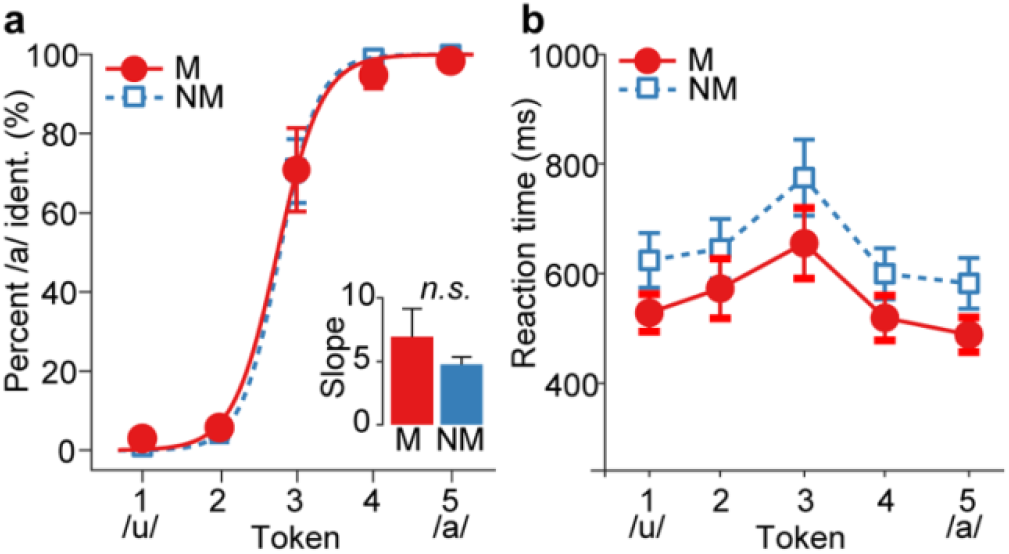
Group performance in speech categorization. Both groups demonstrated similar (a) psychometric function slopes and (b) reaction times when identifying speech sounds along a vowel gradient. More prototypical tokens (those near the extremes of the continuum) were categorized faster than those at the midpoint (i.e., vw 3) indicating slower decisions for category-ambiguous speech sounds. errorbars = ±1 s.e.m.

### Electrophysiological data

To provide comparable measures of response amplitude between brainstem and cortical ERPs, we measured (i) the root mean square (RMS) amplitude of the steady state portion of brainstem FFRs (50 - 150 ms window) and (ii) N1-P2 magnitude of cortical ERPs averaged across speech tokens (i.e., vw 1-5). We also evaluated the latencies of ERP components (i.e., P1, N1, P2) and the N1-P2 interpeak latency (IPL). Latencies reflect neural processing efficiency with prolonged latencies suggesting less efficient neural encoding. As MCI is known to compromise processing speed and efficiency^29,30^, we wanted to evaluate whether musical experience influenced processing efficiency in at-risk individuals.

#### Brainstem FFRs

Figure 3a shows average FFRs and response spectra for Ms and NMs for the prototypical /u/ and /a/ tokens (i.e., vw 1 and vw 5). The strength of brainstem speech encoding, as measured via FFR RMS amplitude, was similar across groups (*F*_1, 11_ = 0.65, *p =* 0.44). However, FFR amplitudes varied with age such that older individuals demonstrated more robust brainstem responses than their younger peers (*F*_1, 11_ = 8.44, *p =* 0.01, 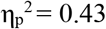; Fig. 3b). Unusually large speech FFRs in cognitively at-risk individuals corroborates previous studies demonstrating overexaggerated brainstem responses in MCI^12^.

**Figure 3.**
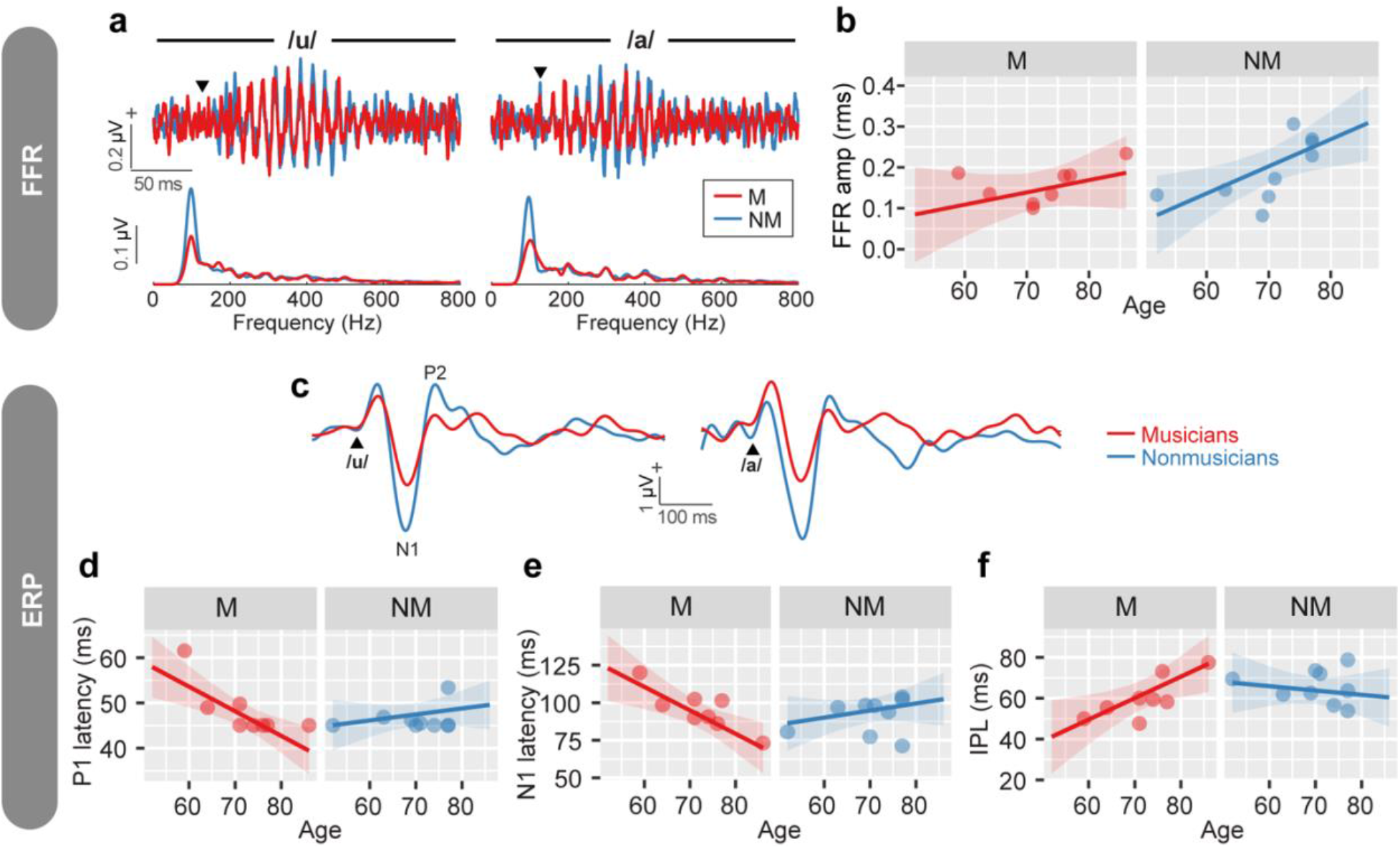
Brainstem FFRs and cortical ERPs are modulated by age and musical experience. Average (a) FFR waveforms and spectra and (c) ERP responses for M and NMs are plotted for the prototypical tokens (i.e., /u/, /a/). (b) FFR amplitudes increase with age but are invariant between groups. (d-e) Cortical P1 and N1 latencies are modulated by age and musical experience. Ms demonstrate more efficient neural encoding with advancing age than their NM peers. (f) NMs showed little change in IPLs with increasing age while Ms demonstrate a gradual increase in IPL. shading = 95% CI

#### Cortical ERPs

Figure 3c shows the average cortical ERPs for Ms and NMs for prototypical /u/ and /a/. N1-P2 amplitude was invariant across groups (*F*_1, 11_ = 0.13, *p =* 0.72). However, stark group and group x age differences were observed for response latencies. Ms demonstrated shorter latencies on average than the NM group for the P1 and N1 waves (P1: *F*_1, 11_ = 12.30, *p =* 0.005, 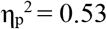, Fig. 3d; N1: *F*_1, 11_ = 10.31, *p =* 0.01, 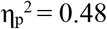, Fig. 3e). Group × age interactions revealed Ms demonstrated decreased P1 and N1 latencies with increasing age while NMs demonstrated little age-related change (P1: *F*_1, 11_ = 12.22, *p =* 0.005, 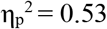, Fig. 3d; N1: *F* = 10.53, *p =* 0.01, 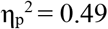, Fig. 3e). Group differences were also noted in N1-P2 IPL (*F*_1, 11_ = 6.42, *p =* 0.03, 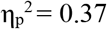, Fig. 3f) with responses being more prolonged in NMs than Ms. However, the group × age interaction was also significant (*F*_1, 11_ = 6.05, *p =* 0.03, 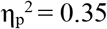; Fig. 3f); whereas IPLs increased with age in Ms, NMs’ ERPs were relatively invariant (though they were longer overall). The prolonged IPL in Ms was driven by the age-related speeding up of N1 latency as neither group nor age differences were noted for P2 latency (group: *F*_1, 11_ = 2.15, *p =* 0.17; age: *F*_1, 11_ = 0.27, *p =* 0.62). Thus, because Ms’ N1 latency decreased with age and no change was observed for P2, the time elapsed between the occurrence of N1 and P2 (i.e., N1-P2 IPL) was prolonged. This is further suggested by the negative correlation between N1 latency and IPL (*r =* −0.83, *p*_adj_ = 0.001). No other significant effects or interactions were found.

### Relations between neural measures, musical experience, and cognitive status

Pairwise Pearson correlations allowed us to determine the degree to which musical experience influenced brainstem and/or cortical speech encoding in individuals at risk for MCI. The analyses between neural measures, musical experience, and MoCA scores revealed cognitive status was predicted by brainstem response amplitude (Fig. 4a); larger speech FFRs were associated with lower (poor) MoCA scores (*r =* −0.60, *p*_adj_ = 0.05). Critically, we found that listeners’ duration of musical experience related to auditory cortical response latency and amplitudes (Fig. 4c, d). More extended musical experience was associated with later P1 latency (*r =* 0.69, *p*_adj_ = 0.02; Fig. 4c). Additionally, musical experience was inversely related to N1-P2 amplitude (*r =* −0.62, *p*_adj_ = 0.05; Fig. 4d) suggesting listeners with less musical experience have abnormally robust cortical speech responses, as observed in more severe forms of MCI and cognitive aging^12^. Though trending in the predicted direction, musical experience did not predict cognitive scores (*r =* 0.42, *p*_adj_ = 0.20; Fig. 4b).

**Figure 4.**
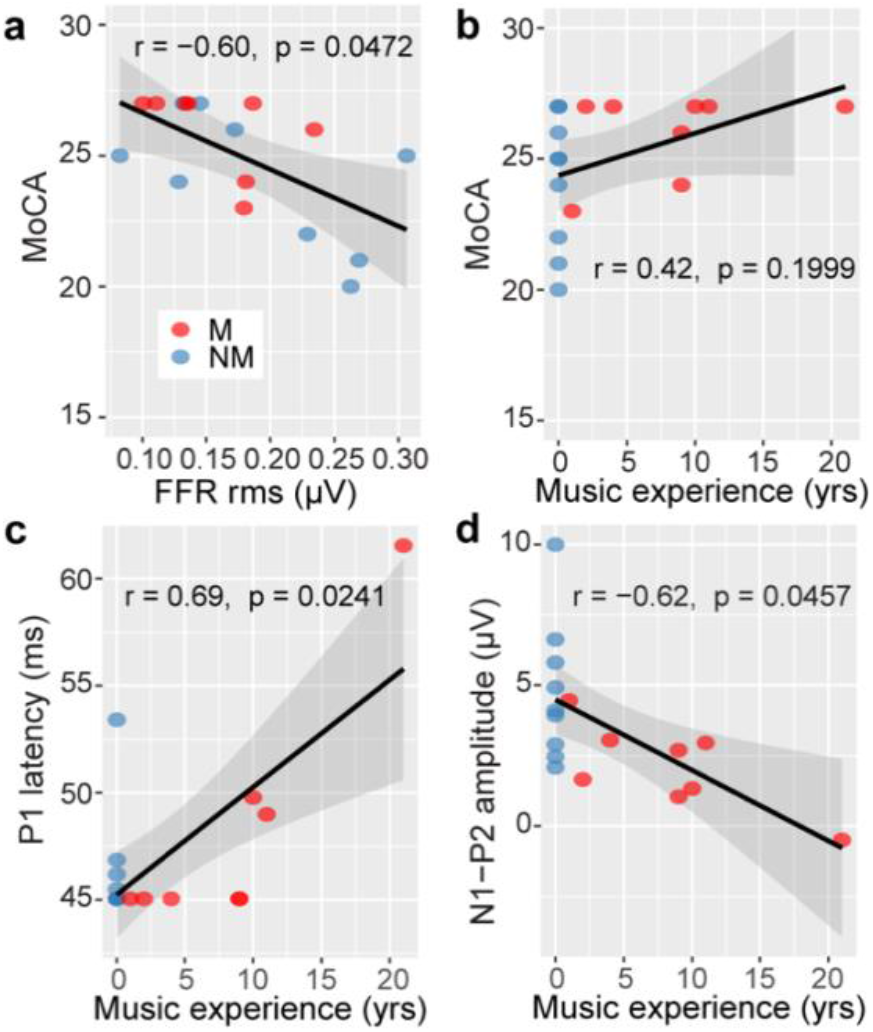
Correlations depicting relationships between MOCA scores, musical experience, and neural measures. (a) Cognitive status is predicted via brainstem FFRs; larger FFRs are indicative of lower (poorer) MOCA scores. (b) Extent of musical experience was not associated with degree of cognitive impairment, *per se*. (c-d) Longer duration of musical experience is associated with later P1 latencies and smaller N1-P2 amplitudes. *p* values reflect FDR adjustments for multiple comparisons. shading = 95% CI

## DISCUSSION

Typical aging degrades neural representations of speech producing weaker encoding and timing delays in auditory brain processing that are further exacerbated by MCI^6–8,12^. By recording speech-evoked potentials via single-channel EEG, we sought to evaluate the neuroplastic effects of prior music experience in older adults with putative MCI. Our data reveal (i) overly robust brainstem FFRs predict severity of cognitive impairment (MoCA scores) and (ii) musical experience offsets neural processing delays in older adults with atypical cognitive aging.

In our sample of individuals with borderline MCI, cognitive function correlated with FFR amplitudes but not cortical responses. Similarly, Bidelman, et al. ^12^ reported that brainstem measures better distinguish older adults with and without MCI than cortical responses alone. More specifically, we found those with poorer MoCA scores—indicative of more severe cognitive impairment—demonstrated overexaggerated FFRs. These findings corroborate previous work suggesting individuals with MCI demonstrate hypersensitive auditory sensory responses across both subcortical and cortical levels of processing^12,13,31,32^. Hypersensitive encoding in MCI could be due to disinhibition in feedback from higher-order brain areas (e.g., prefrontal cortices) which normally regulate early auditory encoding via sensory gating mechanisms^31–33^. Yet, altered feedback from prefrontal cortices would likely influence neural representations within both subcortical and cortical levels which is not observed here. Changes local to brainstem have also been identified in preclinical stages of Alzheimer’s disease^34^. Thus, the overly robust subcortical responses we observed in those with more severe cognitive impairment most likely result from localized changes within the brainstem that precede more obvious stages of cognitive decline. The lack of correlations between cortical ERPs and cognitive function might be due to the mild severity of cognitive deficits in our sample. It is possible that cognitive decline was not yet egregious enough to influence the robustness of auditory processing at the cortical level. Indeed, primary and secondary auditory cortices are not directly influenced by neuropathological changes in cognition until more advanced stages of the disease^35^. Yet, that subcortical enhancements scale with MoCA scores even in a borderline clinical population suggests that brainstem FFRs are highly sensitive to pre-clinical cognitive deficits and may serve as an early biomarker to index cognitive decline prior to traditional neuropsychological tests^12,16^.

Our results also suggest that musicianship might mitigate the normal decline in temporal processing commonly observed in aging adults. Aging results in poorer neural precision and synchrony which is thought to contribute to perceptual temporal processing deficits in older adults, including poor speech-in-noise understanding^5,6,8,25,36,37^; furthermore, cognitive impairment exacerbates these typical age-related declines in processing efficiency^10,13,15^. Here, we show that older adults at risk for MCI but with prior musical experience have faster cortical responses to speech with advancing age than their nonmusician peers who are similarly susceptible to cognitive aging. Critically, the differential effect of age in older musicians vs. nonmusicians is promising in that it demonstrates music engagement later in life might help slow the progression of cognitive decline, particularly in those most at risk. Still, while the correlational nature of our data is provocative, larger scale and/or longitudinal training studies are needed to confirm whether music enrichment programs might truly offer an effective intervention for MCI.

Prior work suggests musical experience enriches neural representations of speech across the auditory hierarchy and improves auditory skills necessary for speech understanding^23,38,39^. The functional, neuroplastic changes due to musical training experience, even when completed early in life, continue to impact auditory processing and speech perception into adulthood^18–20,40^. Training benefits have been shown to limit typical aging effects, resulting in higher fidelity (i.e., amplified and more temporally precise) neural encoding^18,25^. Our results provide further support for notions that musical experience counteracts normal age-related declines in temporal precision by extending those findings to atypical cognitive aging. Furthermore, our findings provide new evidence to suggest brief to moderate musical experience attenuates auditory neural aging effects as several of our participants received only minimal experience (i.e., < 5 years). Recent studies confirm the benefits of short-term musical experience on the neural encoding and perception of normal and even noise-degraded speech in older adults^26,41^.

The earlier cortical ERP latencies in musicians suggest these individuals exhibit more efficient sensory encoding of auditory inputs. P1 and N1 reflect the initial registration of acoustic stimuli within early primary and secondary auditory cortices^42–44^. Because these components dominantly reflect exogenous stimulus properties^43,44^, shorter ERP latencies in musicians imply that musical experience improves initial cortical processing efficiency for complex stimulus features. Seemingly in contrast, our correlational analyses revealed P1 latency increases with degree of musical training. This relationship appears to be driven by the participant with the greatest extent of experience (21 years) who also is the youngest in the M group (Fig. 4c). This result further highlights the influence of age on auditory processing efficiency in older musicians. The age-related increases in IPLs (N1-P2 latency) in those with musical experience appear to result from reduced latencies mainly in the N1 (rather than P2). The lack of group differences in P2 latency might be expected given this wave indexes stimulus classification^45,46^ and we did not find group differences in behavioral categorization. In sum, our ERP latency data suggest that musicians processed speech sounds more efficiently than their nonmusician peers allowing more time for the perceptual decision-making process. Still, RTs were similar (though in the correct direction) between groups so the behavioral relevance of these neural effects remains speculative. For future studies, it would be interesting to evaluate whether changes in cortical IPLs relate to perceived effort in speech perception or participants’ confidence in response selection. For example, musical experience might make speech processing less demanding in cognitively at-risk older adults even if it does not yield strong advantages at the behavioral level^47,48^.

We found no group differences in cortical amplitude measures which suggests latencies may serve as a more sensitive measure of neuroplastic effects in individuals at risk for MCI. This result is consistent with previous studies showing differences in cortical responses between individuals with and without musical experience in the later auditory potentials (i.e., > 100 ms)^23,49–51^. Despite the lack of a group effect, we did find a strong association between degree of musical experience and ERP amplitudes: individuals with more experience demonstrated less robust N1-P2 amplitudes. This result replicates previous work revealing musical experience modulates N1-P2 amplitude^52,53^. Tasks requiring focused attention tend to increase N1 amplitude^43,54^. Thus, decreased N1-P2 in Ms may indicate the task was less cognitively taxing (i.e., requiring less attentional resources) for those with more musical experience. As with the latencies, the reduction in N1-P2 amplitudes suggests that the task might have required less neural activation to produce equivalent behavioral speech categorization and further advances the notion of improved neural efficiency in those with more extensive musical background. That the impact of musical experience is restricted to cortical responses replicates prior work suggesting musical training does not influence the robustness of the steady-state FFR in older adult listeners^18,55^. Furthermore, because age reduces FFR amplitude^6,36,37^, it is possible the neuroplastic benefits of musical experience in subcortical levels of processing, if present in this population, were not robust enough to be detected in our small sample (M: *n =* 8).

Previous studies have separately focused on the impact of MCI or musical experience on auditory processing in older adults. Here, we examined the combined effects of musical experience in individuals at risk for MCI to directly assess the interaction between these positive (experience-dependent; musicianship) and maladaptive (MCI) forms of plasticity. Our results bolster the novel utility of brainstem speech-FFRs as an index of monitoring cognitive decline in individuals with borderline clinical symptomology^12^. They further suggest neurological changes detected in auditory-sensory brainstem processing may serve as an early biomarker for detecting cognitive deficits prior to traditional behavioral metrics^12^. Additionally, our findings provide new evidence that musical engagement might offset typical age-related delays in auditory cortical brain processing resulting in more efficient speech processing in older adults with putative MCI. While our findings are preliminary and await replication in a larger sample, our cross-sectional data opens the door for future training studies examining music intervention as a protective solution for mitigating auditory-perceptual declines due to cognitive aging.

## METHODS

### Participants

Seventeen older adults (*M±SD*, 71.1 ± 8.0 yrs; range: 52-86 yrs) participated in the study. The sample was selected *post hoc* from a larger database of studies examining auditory-cognitive aging, speech processing, and the brain^6,18^ based on two inclusion criteria. First, we selected individuals who might be considered at risk for MCI and early cognitive decline based on lower-than-normal scores on cognitive screening. Cognitive status was assessed by the Montreal Cognitive Assessment (MoCA; published normative cutoff score for normal cognitive function=26)^27^. Participants scoring near or below the normative cutoff for MCI (≤ 27 points; 25 ± 2.3; range: 20-27) were identified as being at-risk or having putative MCI. We included individuals with borderline scores (=27) to (i) assess older adults at risk for MCI and (ii) ensure better *post hoc* matching of groups along other confounding variables (e.g., hearing status, age, education, etc.). While these scores fall within or near the normative range for mild cognitive impairments (19-25.2), they remain above the MoCA normative range for Alzheimer’s Disease or more severe dementia (11.4-21)^27^. Underlying etiology was unknown.

From among those identified with at-risk MoCA scores, we then divided the sample into two groups based on whether listeners had *any* prior musical training. Musicians (Ms; n = 8; 2 female) were defined as amateur instrumentalists who had received at least 1 year of private instruction on their principal instrument. Ms had on average 8.4 ± 6.4 yrs of musical experience. Nonmusicians (NMs; n = 9; 5 female) had no previous musical experience (0.0 ± 0.0 yrs) of any kind in their lifetime. While this definition is perhaps more lax than typical definitions of musicians, we aimed to assess musical experience as a continuous rather than dichotomized trait^56^ so as to not discard individual differences^57^. Given the preliminary nature of this study, treating musical experience as a graded variable also provided us the opportunity to assess if musicianship of any kind, and however minimal, might offer protective effects against MCI.

On average, the groups demonstrated clinically normal hearing (≤ 25 dB HL) based on pure-tone average (PTA) thresholds (i.e., average of 500, 1000, 2000 Hz) bilaterally, and hearing sensitivity did not differ between groups (*t*_15_ = −0.71, *p =* 0.49). Groups were also balanced in age (*t*_15_ = −0.56, *p =* 0.58) and cognitive function as assessed by the MoCA (*t*_15_ = −1.79, *p =* 0.09). Ms reported more years of formal education on average (M: 19.1 ± 3.0 yrs; NM: 14.8 ± 3.6 yrs; *t*_15_ = −2.67, *p =* 0.02); therefore, education was included as a covariate in all subsequent analyses. All participants were right-handed^58^ and native English speakers who reported no history of neuropsychiatric disorders. Participants were compensated for their time and provided written, informed consent in compliance with the Declaration of Helsinki for experiments involving humans and a protocol approved by the Baycrest Centre Research Ethics Committee.

### Stimuli

A synthetic five-step vowel continuum (“vw 1-5”) was constructed so that each 100 ms token would differ minimally acoustically, yet be perceived categorically^46,59^. The first formant (F1) frequency was varied parametrically over five equal steps between 430 and 730 Hz resulting in a stimulus set that spanned a perceptual phonetic continuum from /u/ to /a/. All other stimulus attributes (e.g., fundamental frequency, higher formants) were identical between tokens. For further stimulus details, see Bidelman et al.^46^.

### EEG recording procedures and preprocessing

Data acquisition and response evaluation were similar to previous reports from our laboratory (e.g., ^6,18,46^). Stimuli were presented binaurally at 83 dB SPL via insert earphones (ER-3A, Etymotic Research) with extended acoustic tubing (50 cm) to prevent electromagnetic stimulus artifact from contaminating brainstem responses. During ERP recording, we used a forced-choice procedure in which participants heard 200 randomly ordered tokens and were asked to indicate whether they perceived “u” or “a” via a button press on the keyboard. Following participants’ behavioral response, the interstimulus interval (ISI) was jittered randomly between 400 and 600 ms (20-ms steps, rectangular distribution) to avoid alpha entrainment of the EEG^44, p. 168^ and listeners’ anticipation of their behavioral response. We then collected an additional 2000 passive trials (ISI = 150 ms) to measure the sub-microvolt FFR^60^. Brainstem responses are roughly an order of magnitude smaller than cortical activity, and a larger number of trials is required to obtain a comparable signal to noise ratio for brainstem FFRs and cortical ERPs^61^. During the passive block, participants watched a self-selected movie with subtitles to facilitate a calm yet wakeful state throughout recording.

Continuous EEGs were recorded differentially between an electrode placed on the high forehead at the hairline (~Fpz) referenced to linked mastoids. A third electrode on the mid-forehead served as the common ground. This montage is optimal for recording evoked responses of both subcortical and cortical origin^23,46,62^. Electrode impedances were ≤ 3 kΩ. EEGs were digitized at 20 kHz and bandpass filtered online between 0.05 and 3500 Hz (SynAmps2, Compumedics NeuroScan). Traces were then segmented (cortical ERP: −100-600 ms; brainstem FFR: −40-210 ms), baselined to the pre-stimulus interval, and subsequently averaged in the time domain to obtain ERPs for each condition^63^. Trials exceeding a ±50 μV threshold were rejected as blinks prior to averaging. Grand average evoked responses were then bandpass filtered in different frequency bands to emphasize brainstem (80-2500 Hz) and cortical (1-30 Hz) neural activity, respectively^18,46,61,62^.

### Behavioral data analysis

Psychometric identification functions were created by calculating the proportion of trials identified as a single vowel class (i.e., /a/) for each token (i.e., vw 1-5). Comparing the slope of the psychometric functions between groups allowed us to assess possible differences in the “steepness” (i.e., rate of change) of the categorical speech boundary as a function of musical experience. Behavioral speech labeling speeds, i.e., RTs, were computed separately for each participant as the mean response latency across trials for a given speech token. Following our previous reports on categorical speech perception (e.g., ^18,46^), RTs shorter than 200 ms or exceeding 5500 ms were regarded as implausibly fast responses and lapses of attention respectively and were excluded from analysis.

### Electrophysiological data analysis

To provide comparable measures of response amplitude between brainstem and cortical ERPs, we measured (i) the RMS amplitude of the steady state portion of brainstem FFRs (50 - 150 ms window) and (ii) N1-P2 magnitude of cortical ERPs averaged across speech tokens (i.e., vw 1-5). N1 was identified as the peak negativity between 70 and 120 ms and P2 as the peak positivity between 150 and 250 ms (e.g., ^6,32^). The overall magnitude of the N1-P2 complex, computed as the voltage difference between the two individual waves, was used as a singular index of the total cortical activation across vowel tokens. While other FFR/ERP measures are available in the literature^44,64^, FFR RMS and cortical N1-P2 metrics are advantageous because they both provide a description of the overall amplitude of the evoked response at brainstem and cortical levels using an isomorphic metric. We also evaluated the latencies of the P1 (peak positivity between 45 and 65 ms), N1, and P2 components of the cortical response as well as the N1-P2 IPL.

### Brain-behavior relationships

Pairwise Pearson’s correlations were used to investigate correspondences between subcortical and cortical speech representations (brainstem: RMS amplitude; cortical: P1, N1, P2 latencies and N1-P2 magnitude and IPL), musical experience, and cognitive status. False discovery rate corrections were applied to multiple correlations^65^, and adjusted p-values are reported. This analysis allowed us to determine the degree to which musical experience influenced brainstem and/or cortical speech encoding in individuals at risk for MCI.

### Statistical analyses

We used an analysis of covariance (ANCOVA) on each dependent variable (*lme4*; R)^66,67^ with group (2 levels: M, NM) serving as the between-subjects factor and education, PTA, and age serving as continuous covariates. Following our previous work, responses were averaged across vowel tokens to reduce the dimensionality of the data and directly assess musicianship effects on neural responses^12^. Although groups did not differ in hearing sensitivity, age-related hearing loss is known to alter brainstem and cortical auditory evoked potentials^1,6^. Therefore, we included PTA as a covariate in analyses to control for potential hearing-related differences in neural responses. Cognitive function declines with age^4,30^; therefore, age was also included as a covariate in all analyses. When needed, Tukey-Kramer corrections were used to control Type I error inflation for multiple comparisons. Effect sizes for significant results are given as partial eta-squared 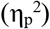.

## Data Availability

The datasets generated and analyzed during the current study are available from the corresponding author on reasonable request.

## ACKNOWLEDGEMENTS

This work was supported by grants from the GRAMMY^®^ Foundation and National Institutes of Health (NIH/NIDCD R01DC016267 and R01DC016267-03S1) awarded to G.M.B. Requests for materials should be addressed to G.M.B.

## AUTHOR CONTRIBUTIONS

G.M.B. designed the experiment; C.N.P. and G.M.B. acquired and analyzed the data, interpreted the results, and contributed to the writing of the manuscript.

